# New insight into plant intramembrane proteases

**DOI:** 10.1101/101204

**Authors:** Małgorzata Adamiec, Lucyna Misztal, Robert Luciński

**Affiliations:** Adam Mickiewicz University, Faculty of Biology, Institute of Experimental Biology, Department of Plant Physiology, ul. Umultowska 89, 61-614 Poznań, Poland.

## Abstract

The process of proteolysis is a factor involved in control of the proper development of the plant and its responses to a changeable environment. Recent research has shown that proteases are not only engaged in quality control and protein turnover processes but also participate in the process which is known as regulated membrane proteolysis (RIP). Four families of integral membrane proteases, belonging to three different classes, have been identified: serine intramembrane proteases known as rhomboid proteases, site-2 proteases belonging to zinc metalloproteases, and two families of aspartic proteases: presenilins and signal peptide peptidases. The studies concerning intramembrane proteases in higher plants are, however, focused on *Arabidopsis thaliana*. The aim of the study was to identify and retrieve protein sequences of intramembrane protease homologs from other higher plant species and perform a detailed analysis of their primary sequences as well as their phylogenetic relations. This approach allows us to indicate several previously undescribed issues which may provide important directions for further research.

## INTRODUCTION

Proteolysis is considered as a crucial factor determining the proper development of the plant and its efficient functioning in variable environmental conditions. Proteases are involved in protein quality control and protein turnover processes. The protein quality control includes the hydrolysis of proteins damaged due to mutations or as a result of plant exposure to stressing environmental conditions, as well as the hydrolysis of proteins synthesized in redundant quantities or sorted to an incorrect cell compartment. Protein turnover, in contrast, comprises hydrolysis of proteins which, in a given spatio-temporal context under non-stressing conditions, become unnecessary.

The results of studies performed in recent years reveal, however, that proteolytic enzymes also participate in signal transduction pathways by releasing membrane-anchored transcription factors and anti-σ factors. The process is known as regulated membrane proteolysis (RIP) and occurs widely in all life forms. RIP was proven to participate in a variety of processes including signal transduction pathways, fatty acid and cholesterol biosynthesis, protein quality control, cell adhesion, regulation of the immune system, sporulation or conjugation (Erez et al. 2009). To date, four families of integral membrane proteases have been identified: rhomboid proteases, site-2 proteases (S2P), presenilins and signal peptide peptidases.

Rhomboid proteases form a large, divergent family which is described in a MEROPS database as the S54 family (membrane-bound serine endopeptidases; Rawlings et al. 2016). They are characterized by presence of from six to eight transmembrane domains and a catalytic dyad formed by serine and histidine residues. The catalytic dyad constitutes the protease active site located inside a chamber formed by transmembrane domains. The catalytically active serine residue occurs in the evolutionary conserved motif GxSx and the catalytically active histidine residue precedes two glycine residues, which allow tight helix packing. The three residues form the strongly conserved motif HxxGxxxG (Lemberg and Freeman 2007). Some experimental data also indicate that the tryptophan-arginine dyad (WR) located in loop1 may play an important role in maintaining rhomboid activity (Lemberg et al. 2005).

The internal differentiation within the rhomboid protease family relating not only to a number of transmembrane segments but also consensus sequences of evolutionarily conserved motifs resulted in proposition of subdivision of the rhomboid family to six different phylogenetic clades: PARL, secretase-A, secretase-B, mixed secretases, iRhoms and mixed inactive homologs.

According to this classification, the PARL type proteases are characterized by the presence of 6 α-helixes forming the core of the catalytic domain and an additional α-helix at their N-terminus. The group of proteins containing, beside the 6 α-helixes of the core, an additional α-helix at their C-terminus are classified as secretase-A type. The characteristic features of this type of proteins are also the conserved WR-motif in the L1 loop and the specific motif adjacent to the catalytically active serine residue GxSxGVYA. Secretase type B covers proteins characterized by the presence of only six transmembrane α-helixes forming the core of the catalytic domain and region of catalytic serine conserved with the motif GxSxxxF. A group of protein sequence elements characteristic for both secretase types was also identified. In the primary structure of these proteins both the conserved WR motif in L1 and the GxSxxxF motif are present. This group of rhomboids was described as mixed secretases. The last clade covers proteolytically inactive proteins containing an additional loop between the first and second transmembrane α-helixes and GPxx instead of the GxSx motif. The clade of inactive proteins was called “iRhoms”. In many organisms, however, other inactive proteolytically rhomboid homologs, which do not contain features typical for “iRhoms” topology, have been identified.

In the *Arabidopsis thaliana* genome 20 gene sequences encoding rhomboid homologs have been indentified (Tripathi and Sowdamini 2006, Lemberg and Freeman 2007b, Garcia-Lorenzo 2006). The protein product of 13 of them contain both evolutionarily conserved motifs with catalytically active amino acids and thus may be considered as proteolytically active. One of these homologs belongs to PARL-type rhomboids 3, others to secretase B-type and nine to 9 mixed-secretase (Lemberg and Freeman 2007). However, the experimental activity confirmation exists only for one of them (Kanaoka et al. 2005). The remaining 7 genes encode the proteins which were described as proteolytically inactive. Among these 4 proteins were described as inactive rhomboid-like homologs and 3 sequences (At2g41160, At3g07950 and At3g56740) were not analyzed in terms of clade membership (Lemberg and Freeman 2007).

The second class of intramembrane proteases comprises site-2-proteases (S2P), zinc metalloproteases occurring widely in living organisms, from bacteria to humans, first identified in humans as protein involved in proteolysis of sterol-regulatory element-binding proteins (SREBPs; Rawson et al. 1997). S2P proteases comprise of at least four hydrophobic regions and their characteristic feature is the presence, in their first transmembrane domain, of the HExxH motif, responsible for zinc ion binding. Two other motifs were also shown to be highly conserved in S2P: GpxxN/S/G located within the second TM and the NxxPxxxxDG located in the third TM. The aspartic acid present in the last motif was shown to be crucial for proteolytic activity of these proteases due to its engagement in zinc ion coordination (Feng et al. 2007). Some S2P proteases also possess PDZ domains, which are known to participate in the interactions between protein molecules forming oligomeric complexes and may play a role in activation of the protease domain (Schuhmann et al. 2012).

Due to its structure S2P has been classified in the MEROPS database as the peptidase M50 family, which is currently divided into two subfamilies. The first one, M50A, consists of S2P human homologs, and the second, M50B, contains homologs of sporulation factor SpoIVFB from *Bacillus subtilis*. These subfamilies are divergent in terms of two structural aspects: catalytic amino acid location and distance between the HEXXH motif and aspartic acid residue. In the M50A subfamilies the catalytic amino acid residues are shifted to the N-terminus in relation to their location in the M50B subfamilies. In turn, the distance between the HEXXH motif and the aspartic acid residue is significantly greater in the M50A subfamily (Rawlings et al. 2016).

There are six genes encoding homologs of S2P in *Arabidopsis thaliana*. The proteolytic activity of four of these proteins was experimentally confirmed, namely: AraSP (Zybailov, et al. 2008), AtEgy1 (Chen et al. 2005), AtEgy2 (Chen et al. 2012), AtEgy3 (Che et al. 2010). The AtEgy3 protein was, in turn, described as proteolytically inactive, since it lacks the HExxH motif and substitution of the aspartic acid in NxxPxxxxDG motif with glutamic acid (Chen et al. 2005).

Presenilins (PSEN) and other aspartic, intramembrane proteases (SPP) are the next intramembrane protease classes, described in the MEROPS database as the A22 peptidase family and divided further into subfamily A, containing homologs of PSEN, and subfamily B, containing homologs of SPP proteases. In terms of topology and catalytic mechanism both classes are very similar – they contain nine transmembrane segments and YDXnLGhGD motif with the two catalytically active aspartates (Erez et al. 2009). What is more, both SPP and PSEN require an additional motif named PAL to maintain proper conformation of their active site (Wang et al. 2006, Tomita et al. 2001). Despite their apparent similarities, SPP and PSEN are characterized by inverse orientation within the membrane (SPP have their C-terminus exposed to the cytosol whereas in the case of PSEN it applies to the N-terminus) and perform the proteolytic cleavage on inversely orientated transmembrane segments. Moreover, PSEN is subject to post-translational processing, which seems not to apply for SPP. Considerable differences also concern functional features of both classes of proteases. SPP act as independently proteases, while PSEN are catalytic subunits of an intramembrane multimeric complex named γ-secretase (Erez et al. 2009). In addition to PSEN and SPP, another group of aspartic intramembrane protease homologs was identified. Since this group of proteins was predicted to share membrane topology with SPP, it was described as SPP-like (SPPL). It is worth noting that while PSEN and SPP proteases were found only in plants and animals, the SPPL were also identified in fungi and archaea (Ponting 2002). The human SPPL2b protein was found to be proteolytically active (Fluhrer et al. 2006, Carpenter et al. 2008).

Two genes encoding presenilins homologs, one encoding SPP protein and five encoding SPPL proteins, were also identified in the *Arabidopsis thaliana* genome. These proteins contain, in their primary structure, all motifs necessary to perform proteolytic cleavage; however, their proteolytic activity has not yet been confirmed.

## METHODS

### Analyzed species

The analysis was performed on nine higher plant species, five of which represent dicots and four monocots. The dicots were represented by *Arabidopsis thaliana*, *Populus trichocarpa*, *Solanum lycopersicum*, *Cucumis melo* and *Vitis vinifera*, and monocots by *Zea mays*, *Brachypodium distachyon*, *Musa accuminata* and *Oryza sativa*.

### Sequence retrieval

The sequences of intramembrane proteases from *Arabidopsis thaliana* were retrieved on the basis of literature research using the TAIR database (Tanya et al. 2015). On the basis of the identified sequences of *Arabidopsis thaliana* proteins the sequences of intramembrane proteins from the remaining 8 species were identified and retrieved using the Aramemnon database (Schwecke et al. 2003). In retrieved sequences the presence of regions with significant homology to one intramembrane proteolytic domain was confirmed using MOTIF, a tool from the GenomeNet platform (http://www.genome.jp/tools/motif/). The sequences for which the proteolytic domain was not detected were excluded from analysis.

### Phylogenetic analysis

The phylogenetic reconstruction was performed using MEGA 6.06. The neighbor-joining method and 1000 bootstrap replications were applied.

### Multisequence alignment

The alignment of sequences within clades and clusters was performed with PRALINE (Simossis and Heringa 2005).

The conserved motifs specific for the given protein family, or clade, were chosen on the basis of literature research. The rhomboid proteases were assigned to the clades on the basis of motifs described by Lemberg and Freeman (2007).

The characteristic motifs were chosen as follows:

- For the PARL clad: sxfsHx_4_Hx_3_Nmx_5_F in the second or third TM, lGaSgax_12_P in the fourth or fifth TM, and aHlgGx_3_G in sixth or seventh TM.
- For secretase B clade: xRx_3_Sx_3_Hx_4_Hx_3_NMx_6_Gx_3_E in the first loop and second TM or in the second loop and third TM, avGxSgvxfx_3_v in the fourth or fifth TM and sfxgHlxGilxG in the sixth or seventh TM.
- For mixed other secretases: WRLx_5_lHx_4_ Hx_3_Nx_7_Gx_3_E in the first loop and second TM or in the second loop and third TM, vGxSgaxfglxg motif in the fourth or fifth TM and aHxgGx_3_G motif in the sixth or seventh TM.

The analysis of above motifs also allowed us to verify the presence of GxSx and HxxGxxxG motifs necessary for proteolytic activity of the proteins.

For characteristic S2P proteins HExxH, AGpxxN/S/G, NxxPxxxxDG motifs were chosen (Feng et al. 2007).

The conserved motifs specific for the aspartic proteases were chosen on the basis of literature research (Erez et al. 2009, Beher et al. 2006, Tomita et al. 2001). The characteristic motifs were identical for presenilins, signal peptide peptidases and signal peptide peptidase like proteins and were chosen as follows: x_5_YDx_5_, x_5_LGxGDx_5_, and x_5_PALpx_5_.

The consensus sequences for analyzed motifs were derived with WebLogo (Crooks et al. 2004).

## RESULTS

In total, 293 protein sequences from nine higher plant species were analyzed (for details see Supplementary material 1). The vast majority of these sequences (164) belong to rhomboid homologs and 48 to site-2-proteases homologs. The aspartic proteases were represented by 81 sequences and among them 18 were described as presenilin homologs, 15 as SPP homologs and 48 as SPPL homologs (for details see Supplementary material 1).

## RHOMBOID HOMOLOGS

The rhomboid homologs were the most numerous among analyzed groups of proteins. The number of sequences in a single species ranged from 14 to 21 (for details see Table 1).

**Table 1.** Number of proteolytically active and inactive proteins assigned to a given clade for every species analyzed.

|  | MONOCOTS |  |  |  | DICOTS |  |  |  |  |
| --- | --- | --- | --- | --- | --- | --- | --- | --- | --- |
|  | *Zea mays* | *Brachypodium distachyon* | *Musa accuminata* | *Oryza sativa* | *Populus trichocarpa* | *Solanum lycopersicum* | *Cucumis melo* | *Vitis vinifera* | *Arabidopsis thaliana* |
| **total** | **21** | **21** | **20** | **18** | **20** | **15** | **14** | **4** | **20** |
| **active** | **12** | **13** | **14** | **13** | **14** | **11** | **11** | **11** | **13** |
| PARL | 0 | 0 | 1 | 1 | 1 | 1 | 1 | 1 | 1 |
| SB | 4 | 4 | 2 | 3 | 4 | 4 | 3 | 3 | 3 |
| MS | 8 | 9 | 11 | 9 | 9 | 6 | 7 | 7 | 9 |
| **inactive** | **9** | **8** | **6** | **5** | **6** | **4** | **3** | **4** | **7** |
| iPARL | 1 | 1 | 0 | 0 | 0 | 0 | 0 | 0 | 2 |
| iSB | 0 | 1 | 0 | 0 | 0 | 0 | 0 | 0 | 0 |
| iMS | 1 | 1 | 1 | 0 | 2 | 2 | 1 | 2 | 2 |
| others | 7 | 5 | 5 | 5 | 4 | 2 | 2 | 2 | 3 |

Among 164 identified sequences, 112 contain both GxSx and HxxGxxxG motifs. Only in a few individual cases, mostly in PARL homologs, glycine residues in HxxGxxxG motif were substituted with alanine residue. Since the alanine residue, like glycine, is small and allows tight packing of the α-helix, we assume that this substitution has little effect on proteolytic activity of the enzyme. The remaining 45 sequences lack at least one of the amino acids that constitute the catalytically active dyad and thus were considered as proteolytically inactive.

All sequences, regardless of their potential proteolytic activity, were analyzed in terms of the presence of motifs characteristic for given rhomboid clades (for details see methods) and phylogenetic identity. The proteins were assigned to 5 different clades (for details see Fig. 1)

**Fig. 1.**
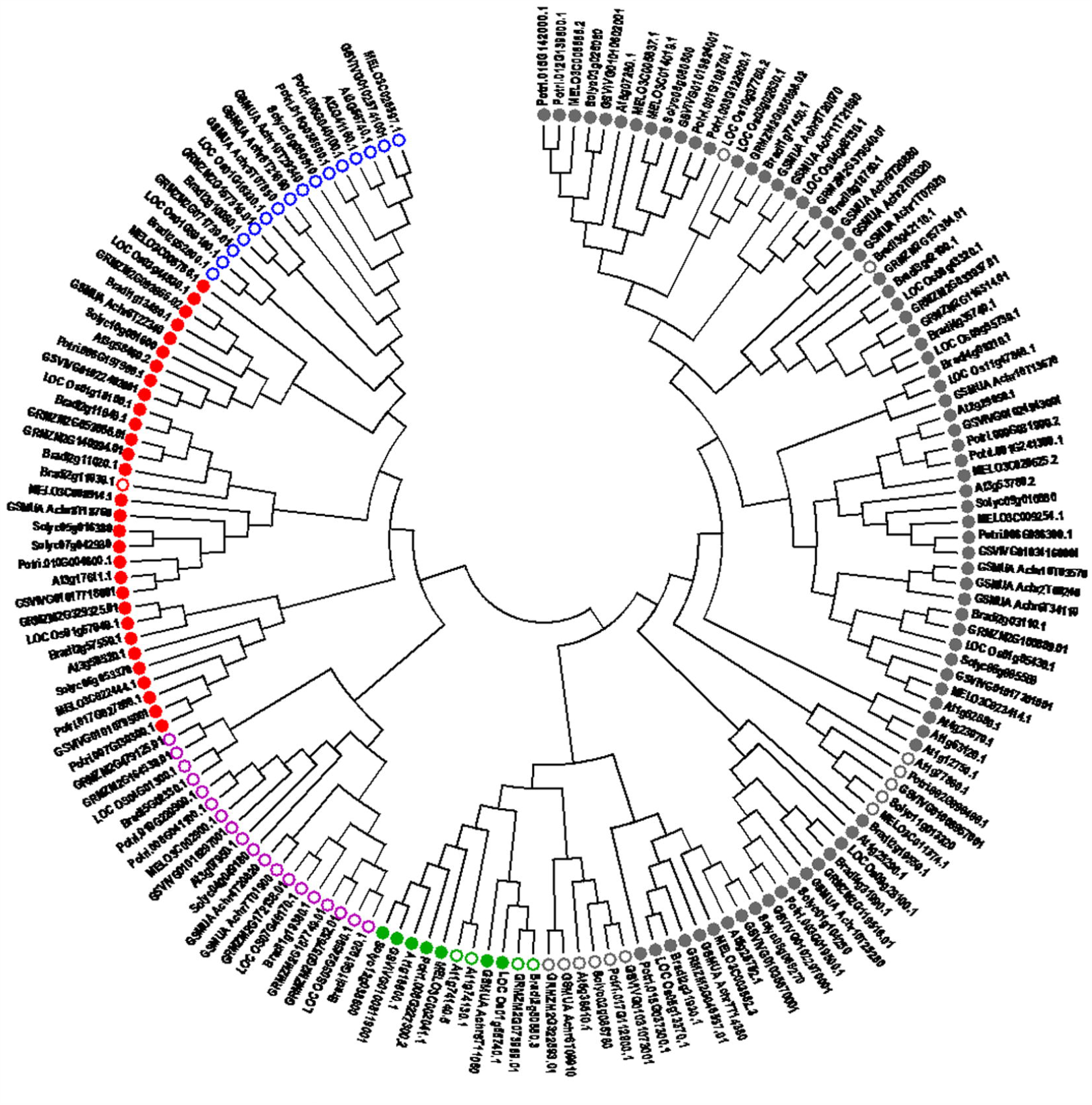
Phylogenetic tree of rhomboid homologs. Potential proteases are marked with a circle and proteins unable to perform proteolysis are marked with a circumference. The rhomboid homolog clades were marked as followed: mixed others (MO) – grey, PARL – green, the Fx – RL purple, SB – red Y-RL – blue.

The analysis revealed 11 sequences containing motifs characteristic for the PARL clade (for motif consensus see Fig. 2). The average identity of these sequences was 43% and identity of their rhomboid domains was 56%. Within this group seven proteins, each belonging to different species, were described as potential proteases since they contain both catalytically active amino acids (for details see Fig. 1). The 4 remaining sequences, two of which belonged to *Arabidopsis thaliana*, one to *Zea mays* and one to *Brachypodium distachyon*, were assigned as proteolytically inactive and described as iPARL proteins. In sequences from *Zea mays* (GRMZM2G073959.01) and *Brachypodium distachyon* (Bradi2g50680.3) proteolytic activity was disabled due to substitution of catalytically active histidine with proline.

**Fig. 2.**
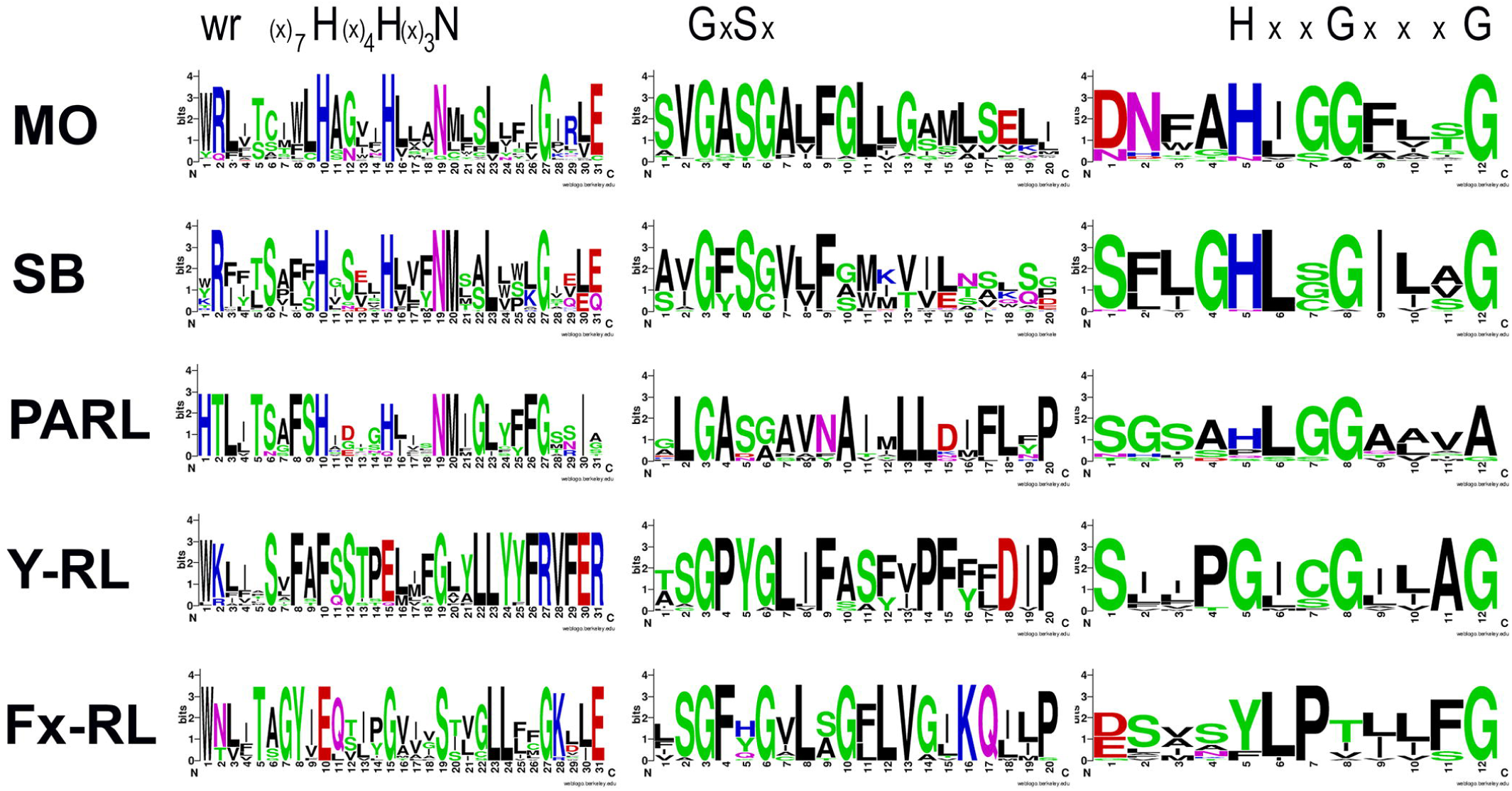
Consensus sequences of motifs characteristic for rhomboid homologs in particular clades.

In *Arabidopsis thaliana* sequences both amino acids that constitute a catalytic dyad were substituted. The serine was substituted with asparagine in At1g74130 or aspartic acid in At1g74140 and histidine was substituted with proline in At1g74130 or serine in At1g74140. It is important to note that 10 proteins belonging to the PARL group were predicted to have 6 transmembrane domains (for details see Supplementary material 1) and for sequence from *Solanum lycopersicum* (Solyc12g038600) only 5 transmembrane domains were predicted. This result is inconsistent with the Lemberg and Freeman (2007) assumption that assignation to clades should be based on the number of transmembrane domains. The results are, however, supported by other publications in which six transmembrane domains were predicted for PARL type proteins from yeast (Rbd1) and *T. gondii* (ROM6; McQuibban et al. 2003, Dowse et al. 2005).

The 31 sequences were assigned to the secretase B clade. The average sequence conservation within these group was 34% and the average identity of their rhomboid domains was 44%. Within this group 14 sequences represented monocots and 17 sequences represented dicots. The number of sequences identified in a single species ranged from 5 to 2 proteins. Among these proteins 30 were described as catalytically active B-type secretases (SB) and only one from *Brachypodium distachyon* (Bradi2g11030.1), due to substitution of catalytically active histidine by asparagine, was classified as inactive (iSB). The predicted number of transmembrane domains within this group was diverse and ranged from 3 to 7. Also in this case, as for PARL-type proteases, the number of predicted transmembrane domains is inconsistent with those proposed by Lemberg and Freeman (2007).

The relatively low sequence identity and divergent number of transmembrane domains within this group prompted us to divide this clade into 3 subclades (for details see Fig. 3). The first subclade consisted of 9 proteins, described as proteolytically active. These proteins shared high identity in all three analyzed motifs (for details see Fig. 4).The most distinctive feature within this group was, however, the GYSC motif containing the active serine residue. The overall sequence identity within this group was 81% and the rhomboid domain average identity was 87%. These proteins were all predicted to possess 7 transmembrane domains; thus we named this group SB7. Among the 9 proteins forming the SB7 subcluster 6 sequences belonged to dicots (2 for *Populus trichocarpa* and one for remaining species) and 3 to monocots (one for *Zea mays*, *Brachypodium distachyon* and *Oryza sativa*).

**Fig. 3.**
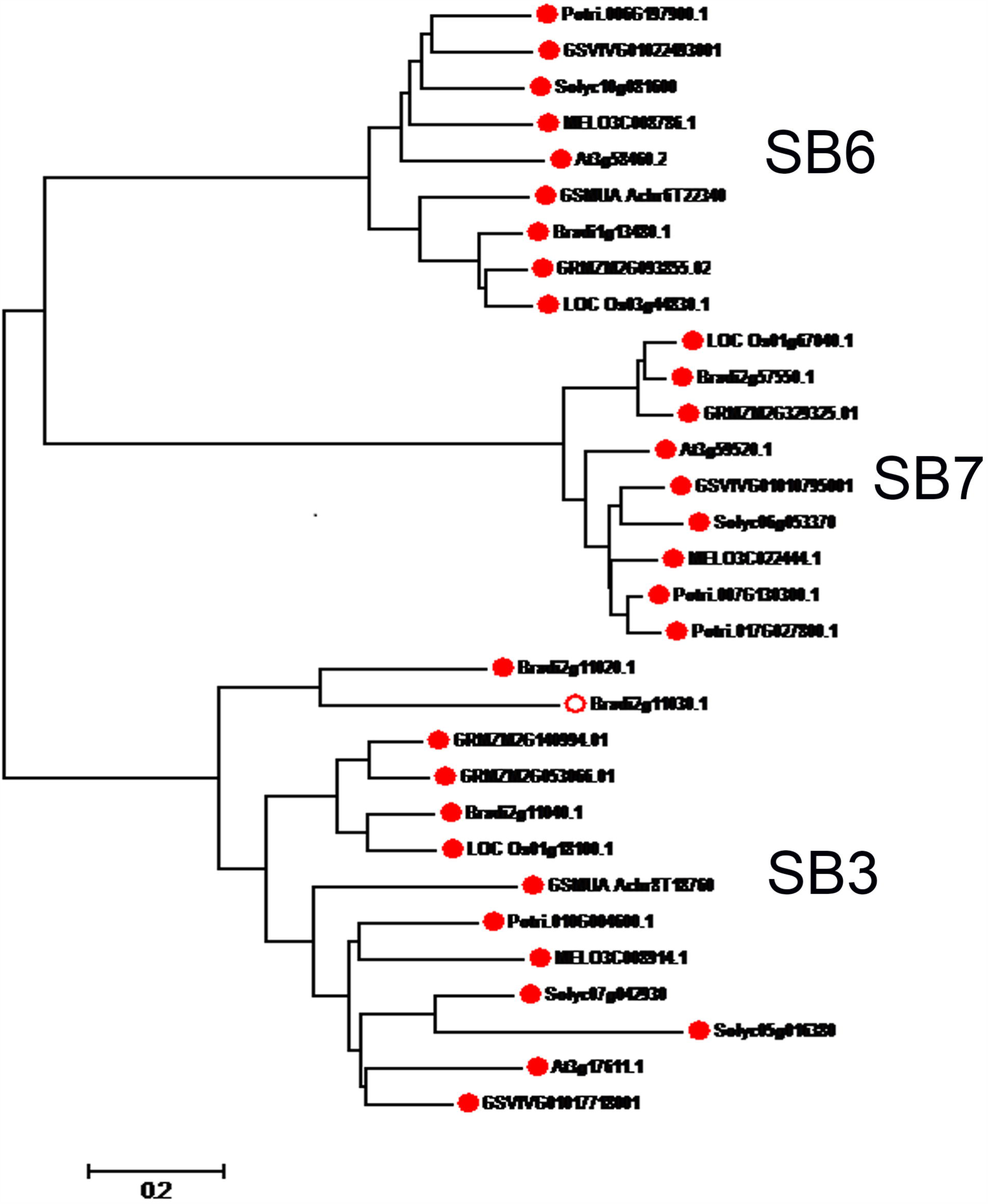
Phylogenetic tree of secretase B homologs with marked division to SB7, SB6 and SB3 subclusters. The potential proteases were marked with a circle and proteins unable to perform proteolytic were marked with a circumference.

**Fig. 4.**
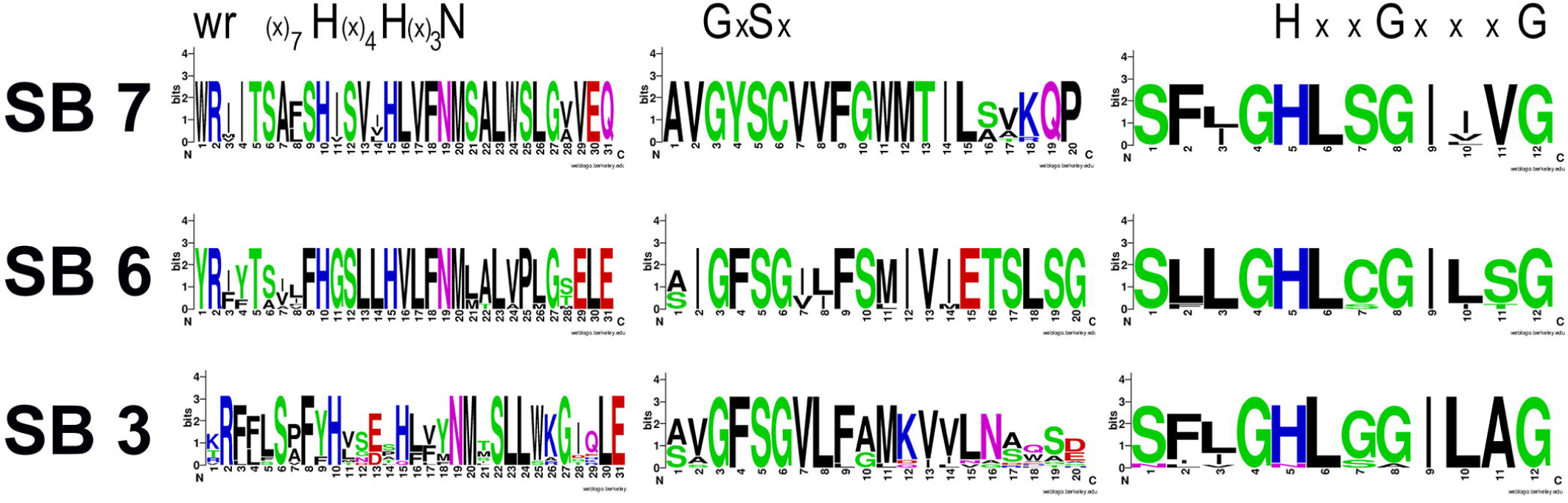
Consensus sequences of motifs characteristic for rhomboid homologs in particular SB subclades.

Also the second subcluster consisted of 9 potential proteases, one for every species analyzed. In every sequence within this subcluster GFSGxxFS and GHLcGILsG motifs were present around catalytically active amino acids (for details see Fig. 4). The average conservation of these sequences was 68%, and the average rhomboid domain identity was 80%. The subcluster was named SB6 since all proteins within this group were predicted to have 6 transmembrane domains.

In every SB6 protease besides the rhomboid domain also a UBA domain was detected. The domain is present in many proteins involved in the ubiquitin/proteasome pathway (Mueller and Feigon 2002).

The third subcluster contains 13 proteins sharing GFSGVL and GHLggILAG motifs in regions responsible for catalytic activity(for details see Fig. 4) The average sequence identity within this cluster was 47%, and the average domain identity was 68%. In all sequences three transmembrane domains were predicted; thus we named this group SB3. In 11 sequences belonging to the SB3 subcluster a zf-RanBP domain was detected. The motif was found in proteins involved in GDP-dependent protein transport and was proven to bind GTP and GDP-bound protein (Yassen and Blobel 1999, Steggerda and Paschal 2002). Among 13 sequences constituting the SB3 subcluster 3 belonged to *Brachypodium distachyon*, two to *Solanum lycopersicum* and *Musa accuminata*, and the remaining species were represented by a single sequence.

The vast group of identified sequences (87 sequences) was assigned to the mixed other secretases clade (for details see Fig. 1). Most of them (75 sequences) were described as proteolytically active. The proteins were predicted to have a divergent number of transmembrane domains, which ranged from 4 to 7. The protein conservation within this group was 40%, and the identity of their rhomboid domains was 51%.

A considerable number of inactive rhomboid homologs (32) were not assigned to any clade. Surprisingly, within this sequence two groups sharing common motifs in analyzed regions were identified.

The first group contained 16 proteins with average sequence conservation 61% and average domain identity 70%. In every sequence within this group catalytically active serine was substituted with tyrosine; thus we described this group as Y-type rhomboid like proteins (Y-RL). In Y-RL proteins the motif GxSx was replaced by the highly conserved GPYG motif and the catalytically active histidine was substituted by a glycine residue in all sequences (for details see Fig. 2). The GPYG motif is similar to the GPxx motif, which was described as characteristic for iRhoms (Lemberg and Freeman 2007); however, our analysis indicates that the Y-RL proteins lack other features characteristic for iRhoms such as an characteristic long loop between the first and second transmembrane domain. For all Y-RL proteins, 5 transmembrane domains were predicted. Thirteen Y-RL proteins share common topology characterized by presence of 3 different proteins domains: DER1, rhomboid and UBA. In the remaining 3 sequences (GRMZM2G157316, GSMUA_Achr5T07850 and GSMUA_Achr8T21690) only rhomboid and UBA domains were present. The Der1 domain was identified as crucial for ER-associated protein degradation in *Saccharomyces cerevisiae* (Knop et al. 1996). Also the UBA domain is present in many proteins involved in the ubiquitin/proteasome pathway (Mueller and Feigon 2002). The presence of both these domains in the studied proteins suggests their involvement in protein degradation. On the other hand, however, the UBA domains are also quite common in cell signaling engaging kinases and proteins involved in DNA excision repair (Mueller and Feigon 2002).

The second subcluster of rhomboid like proteins consists of 19 proteins, whose average sequence conservation equals 60% and average domain identity 64%. The number of transmembrane domains predicted for these sequences ranged from 4 to 6. Within this group of proteins in place of catalytically active serine four different amino acids were present; however, the amino acid was always preceded by phenylalanine forming the GFxG motif (for details see Fig. 2) The catalytically active histidine was substituted mostly by tyrosine; however, in 3 sequences exchange for phenylalanine occurred and in one sequence deletion of this region was observed (for details see Supplementary material 1). Also the amino acids surrounding the residue corresponding to active histidine were relatively weakly conserved; thus the consensus of the motif was ylPxxxfG (for details see Fig. 2). We named this group of proteins Fx type rhomboid like (Fx-RL).

## SITE 2-PROTEASE HOMOLOGS

In the analyzed species 48 sequences containing the M50 domain or its fragment were identified. 27 of these sequences were found in monocots and 28 in dicots (for details see Fig. 5). Detailed analysis of the primary structure of these 48 proteins revealed that 32 of them contain all three conserved motifs important for their catalytic activity. These proteins were considered as proteolytically active. In the remaining 16 sequences at least one of the crucial motifs was missing, and we describe these sequences as proteolytically inactive. The phylogenetic analyses of all of 48 sequences revealed the presence of 5 different clusters (for details see Figure 5). The average sequence conservation within these clusters ranged from 76% to 37%. Surprisingly, the most conserved cluster (76% average sequence identity) consisted of 9 proteins described as inactive. Each sequence within the cluster was derived from different species; thus for every species analyzed only one sequence was present. Since *Arabidopsis thaliana* was represented within this group by AtEgy3 protein, we named this group Egy3-like proteins (Egy3-L). The Egy3-like proteins contained only the short C-terminal fragment of the M50 domain deprived of HExxH and AGpxxN/S/G motifs. The fragment was approximately 36 aa in length and contained the NxxPxxxxDG motif; however, in all Egy3-L proteins the substitution of aspartic acid for glutamic acid occurred (for details see Figure 6). For every sequence within this group 6 transmembrane domains were predicted and the average identity of the fragment of the M50 domain was 95%.

**Fig. 5.**
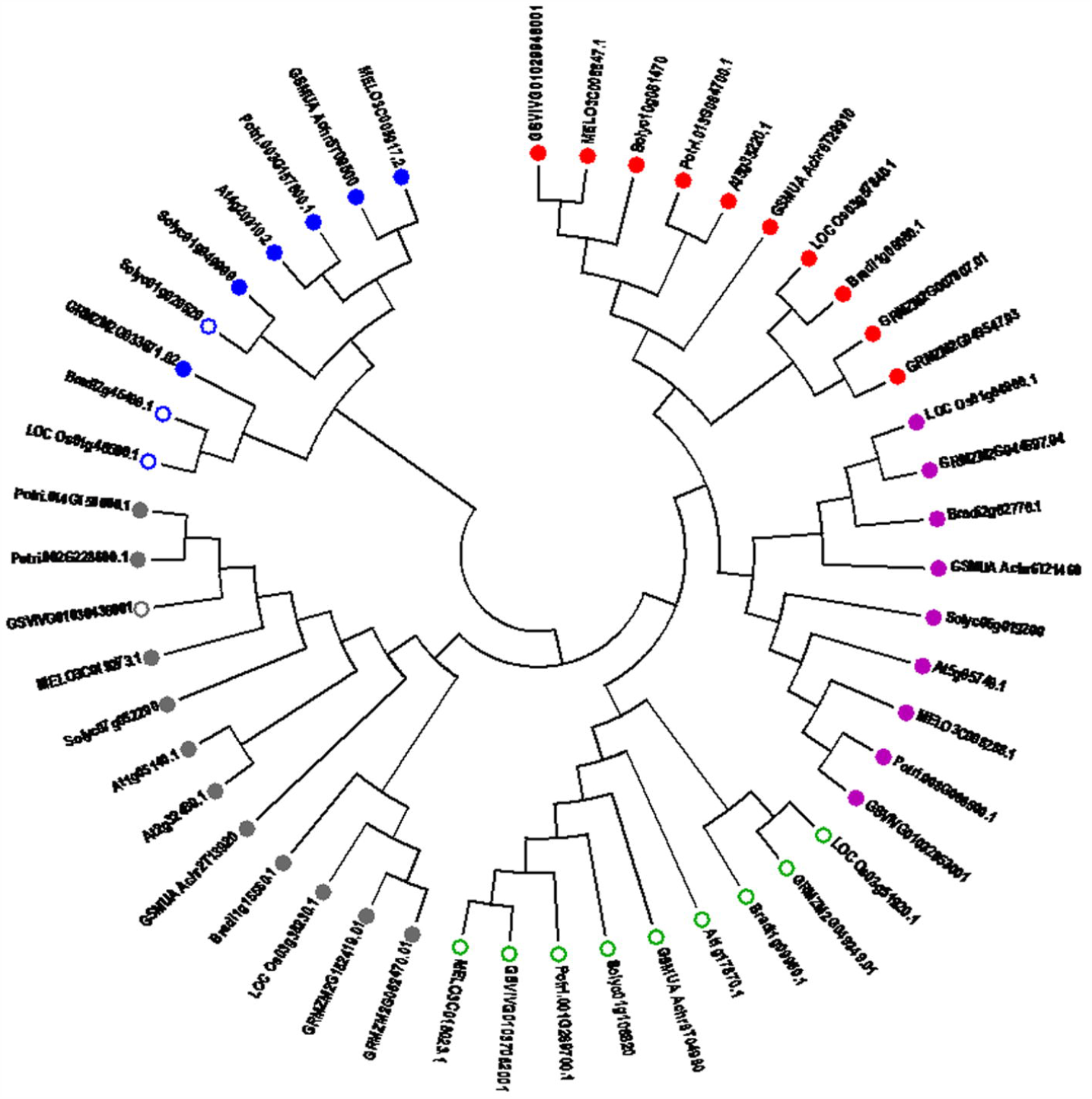
Phylogenetic tree of site-2-proteases homologs. The potential proteases were marked with a circle and proteins unable to perform proteolysis were marked with a circumference. The S2P homologs clades were marked as followed: Egy1-L –red, Egy2-L – purple, Egy3-L green, AraSP-L – grey, Egy4-L – blue.

**Fig. 6.**
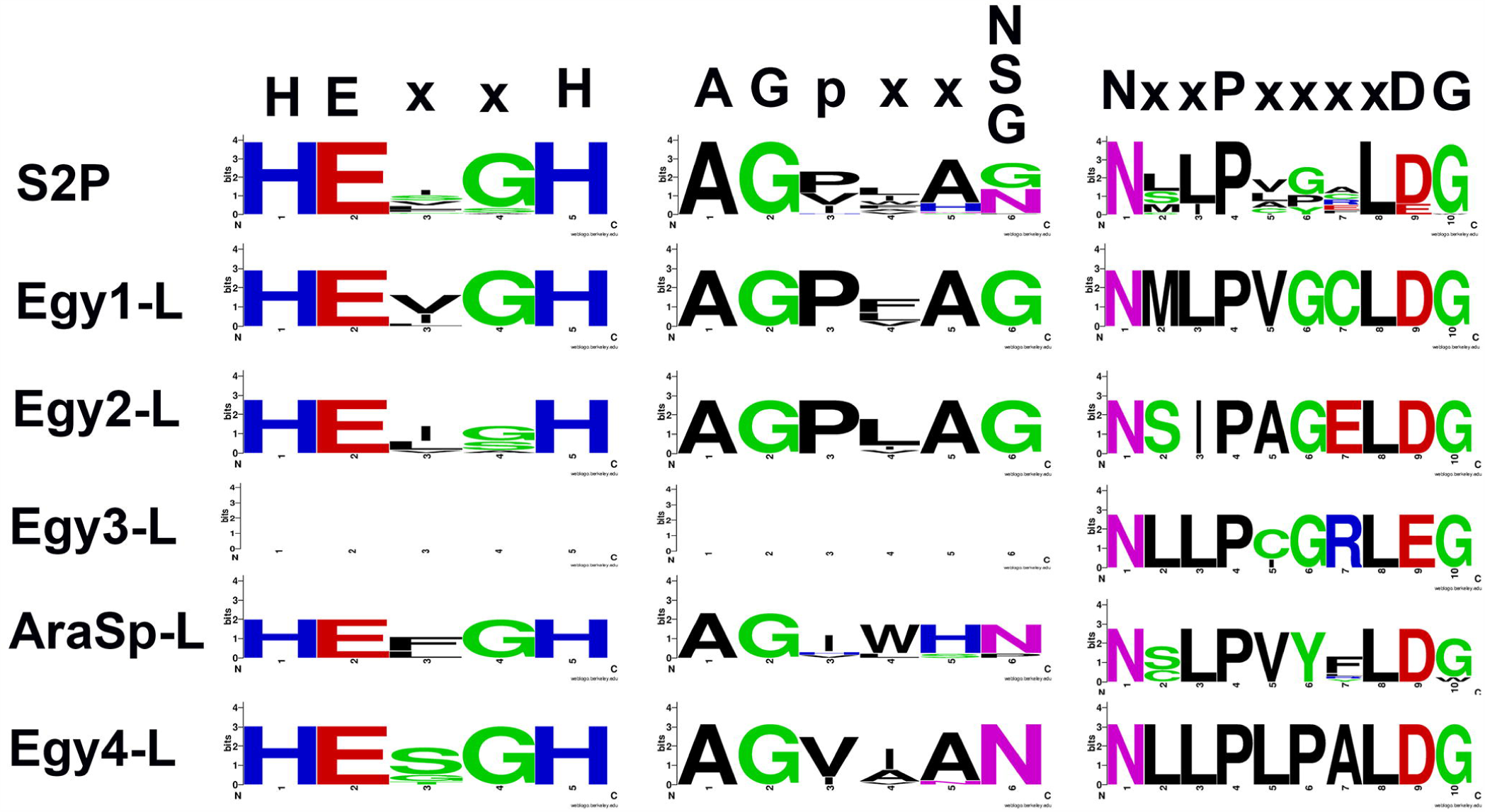
Consensus sequences of motifs characteristic for site-2- proteases. In the first line, named S2P, the consensus for all 48 sequences of S2P homologs is shown. The lines below show consensus sequences of motifs for particular clades.

The second cluster consisted of 10 proteins, whose average sequence identity was 71%. Two of these sequences were derived from *Zea mays* and each of the remaining 8 species was represented by a single sequence. The proteolytic activity of, belonging to this group, AtEgy1 protein (At5g35220) was experimentally confirmed and all remaining proteins possess the 3 conserved motifs crucial for their proteolytic activity (for details see Figure 7) and were described as active proteases. The average domain identity within the Egy1-like cluster was 87%. The proteins within Egy1-L were mostly predicted to possess 7 transmembrane domains, with the exception of sequences from *Oryza sativa* (LOC_Os03g57840.1) and *Brachypodium distachyon* (Bradi1g06080.1), where 8 transmembrane domains were predicted.

**Fig. 7.**
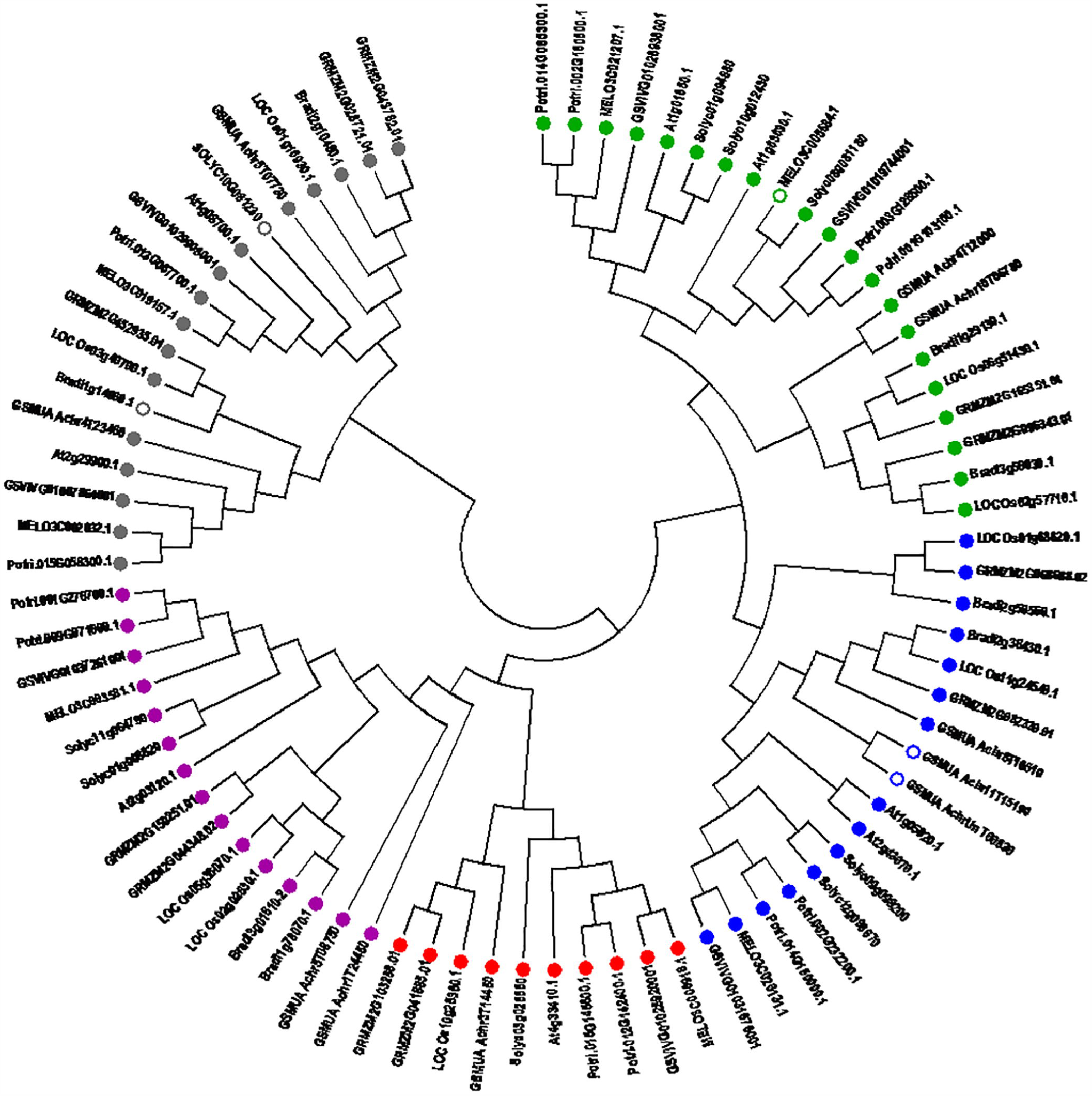
Phylogenetic tree of aspartic proteases homologs. The potential proteases were marked with a circle and proteins unable to perform proteolysis were marked with a circumference. The aspartic proteases homolog clades were marked as followed: SPPL-A – green, SPPL-B – blue, SPPL1 – red, SPP – purple, PSEN – grey.

The third cluster contains 9 proteins with 65% average sequence conservation and 79% domain identity. Within this group one sequence was present for every species analyzed. Two sequences were described as proteolytically inactive. The sequence from *Musa accuminata* (GSMUA_Achr6T21460) lacked AGpxxN/S/G, which was described as essential for active site stabilization (Feng et al. 2007), and in the sequence from *Solanum lycopersicum* (Solyc06g019200) deletion of HExxH and AGpxxN/S/G occurred (for details see supplementary material Alignment). The remaining 7 sequences possess all 3 conserved motifs and were describe as proteases. Proteolytic activity of AtEgy2 (At5g05740), belonging to this cluster was, previously, experimentally confirmed (Chen et al. 2012). For most proteins within Egy2-like cluster 7 transmembrane domains were predicted, except GRMZM2G044697.04 from *Zea mays*, where the number of transmembrane domains was determined as 6.

The fourth cluster consists of 12 proteins whose average sequence conservation equal 64% and average domain identity 70%. Within this group two sequences represented *Arabidopsis thaliana*, *Zea mays* and *Populus trichocarpa* and the remaining 6 species were represented by a single sequence. All sequences within this cluster are potentially able to perform proteolytic cleavage. The proteolytic activity of AraSP (belonging to this group) has been confirmed experimentally (Zybailov et al. 2008). All proteins within the AraSP-like cluster were predicted to have 4 transmembrane domains and a PDZ_2 domain located inside the M50 domain. The PDZ_2 domain is known to be responsible for protein-protein interactions and organization of signaling complexes within the cell membranes (Ranganathan and Ross 1997, Cowburn 1997). The average M50 domain identity within this group was 70%.

The last, weakly conserved group of proteins (37% average sequence identity) contains 8 sequences from 7 species – two *Solanum lycopersicum* sequences and one from *Brachypodium distachyon*, *Musa acuminate*, *Oryza sativa*, *Arabidopsis thaliana*, *Populus trichocarpa* and *Cucumis melo*. The sequence from *Arabidopsis thaliana* included in the cluster was a protein encoded by the At4g20310 gene named by us as Egy4. Within the Egy4-like cluster only 4 sequences have all three conserved motifs and were considered as potentially active and in 7 proteins the PDZ_2 domain was identified (for details see supplementary material). The average domain identity within this cluster was 40%.

## ASPARTIC INTRAMEMBRANE PROTEASES HOMOLOGS

In nine analyzed higher plant species 81 sequences encoding aspartic proteases or their homologs were found. The peptidase _A22B domain, characteristic for SPP and SPPL proteins, was present in 63 of these sequences and in 18 the presenilin domain was identified (for details see table 5). Most of the analyzed sequences (76 of 81) contain all motifs required for their proteolytic activity; thus they may be considered as proteolytically active (for details see Fig. 7). The phylogenetic analysis revealed the presence of 5 clusters: one containing sequences with the presenilin domain, one containing homologs of AtSPP proteins, and three containing AtSPPL homologs.

The most conserved proteins constituted a cluster of 15 proteins sharing high sequence identity within the AtSPP protein. The average conservation within the SPP cluster was 81% and average domain identity was 85%. All proteins within this cluster contain strictly conserved motifs necessary for proteolytic activity (for details see Fig. 8) and were described as potentially active. The majority of proteins were predicted to have nine transmembrane regions; however, for 3 sequences, one from *Cucumis melo* (MELO3C003581.1), one from *Musa accuminata* (GSMUA_Achr1T24450) and one from *Zea mays* (GRMZM2G044348.02), 8 transmembrane domains were predicted.

**Fig. 8.**
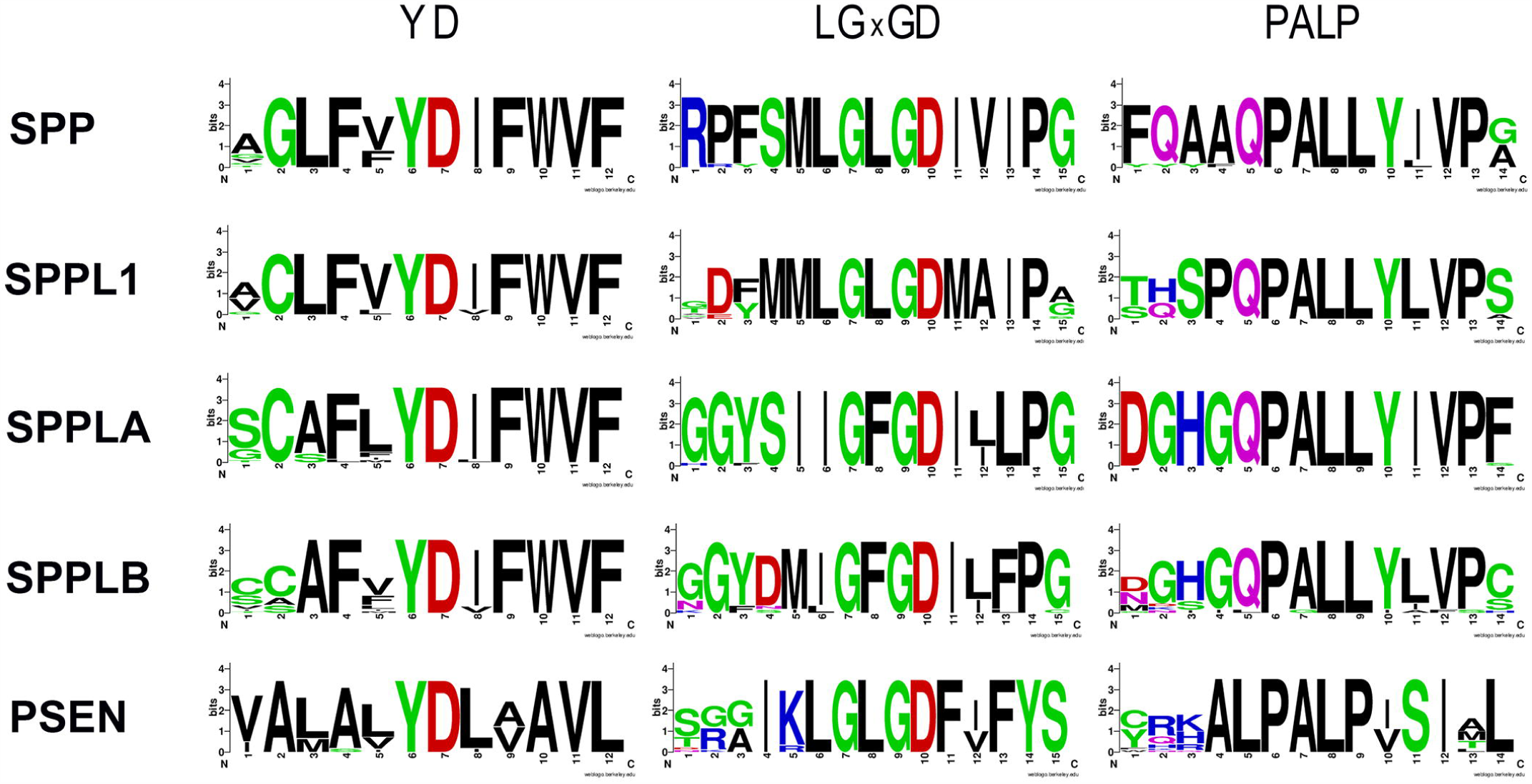
Consensus sequences of motifs characteristic for aspartic protease homologs in particular clades.

Another cluster sharing high sequence identity was formed by 10 proteins, including AtSPPL1 protein. The protein conservation within this group was 78% and the domain identity was 80%. All proteins were potentially able to perform proteolytic cleavage (see Fig. 7) and were characterized by high identity of crucial motifs (for details see Fig. 8). However, only for 4 of these sequences nine transmembrane regions were predicted. The remaining 6 sequences were predicted to have 8 transmembrane domains (for details see supplementary material). The cluster was described as SPPL1.

The third cluster of highly conserved sequences, named SPPL-A, was formed by 21 proteins whose average sequence identity was 74%. The average peptidase_A22B domain identity within this group was 81%. Only one protein within this cluster was predicted as proteolytically inactive (MELO3C005684.1) since it lacked both LGxGD and PALP motifs. In all remaining proteins the motifs were strictly conserved (for details see Fig. 8). This group of proteins was also characterized by presence of protease-associated (PA) in 20 of 21 sequences (except GRMZM2g165351 from *Zea mays*). The average identity of this domain was 69%. The PA domain was previously described in other protease families such as subtilases and zinc metalloproteases, and it was suggested to form an active site covering lid structure (Luo and Hofmann 2001). Proteins within this cluster were generally predicted to have 9 transmembrane domains; however, for one sequence from *Cucumis melo* (MELO3C005684.1) and one from *Zea mays* (GRMZM2G165351.03) 6 and 7 transmembrane domains, respectively, were predicted.

The sequence conservation in the two remaining clusters was significantly lower. The last cluster containing SPPL homologs (SPPL-B) was characterized by 56% average sequence identity and 65% average peptidase_A22B domain identity. The cluster was formed by 17 proteins and included SPPL3 and SPPL5 proteins from *Arabidopsis thaliana*. Despite relatively low sequence conservation, the motifs necessary to perform the proteolytic cleavage and their proximity were highly conserved (for details see Fig. 8). Only two sequences within this group, both derived from *Musa accuminata*, were considered as proteolytically inactive (for details see Fig. 7). GSMUA_Achr11T15190 lacked the LGxGD motif and GSMUA_AchrUn_T00830 was deprived of both LGxGD and PALP motifs. In the remaining sequences the motifs crucial for proteolytic activity were strictly conserved. The sequences within this cluster contain, apart from the peptidase_A22B domain, also the PA domain. The exceptions were one sequence from *Solanum lycopersicum* (Solyc09g098200) and one from *Musa accuminata* (GSMUA_Achr8T16510) where the domain was absent. The average identity of the PA domain within this group was 56%. In the vast majority of sequences nine transmembrane regions were predicted; there were, however, four exceptions. For two sequences, one from *Solanum lycopersicum* (Solyc09g098200) and one from *Musa accuminata* (GSMUA_AchrUn_T00830), 7 transmembrane domains were predicted; GSMUA_Achr11T15190, was predicted to have 8 transmembrane domains and LOC_Os01g68620.1 from *Oryza sativa* to have 10 transmembrane domains.

Surprisingly, the least conserved cluster contains presenilin homologs. The cluster was formed from 18 sequences with average sequence conservation of 52% and average domain identity of 53%. In the vast majority of sequences within this group the motifs essential for proteolytic activity and their proximity were highly conserved (for details see Fig. 8), and only two proteins were described as proteolytically inactive (for details see Fig. 7). The sequence from *Solanum lycopersicum* (Solyc10g081230) was deprived of both catalytically active aspartic residues and the sequence from *Brachypodium distachyon* lacked LgxGD and PALP motifs. For most sequences within the presenilin cluster nine transmembrane domains were predicted, but for 5 sequences the number was smaller. For the sequence Solyc10g081230 (from *Solanum lycopersicum*) only 3 transmembrane domains were predicted. The sequence was however deprived of a major part of the peptidase_A22B domain and described as proteolytically inactive due to absence of both crucial active residues. The two sequences from *Musa accuminata* (GSMUA_Achr4T23460 and GSMUA_Achr5T07730), both described as potential proteases, were predicted to have six transmembrane domains. The LOC_Os01g16930.1 sequence from *Oryza sativa* and MELO3C019157.1 from *Cucumis melo*, both potentially able to perform proteolytic cleavage, were predicted to have, respectively, 7 and 8 transmembrane domains.

## CONCLUSIONS

Knowledge concerning intramembrane proteases in the higher plants is, generally, very limited. However, the performed data mining and literature research together with sequence and phylogenetic analysis allow deeper insight into available information and highlight several issues which may provide important directions for further research.

In higher plant rhomboid homologs the number of transmembrane domains weakly corresponds with the presence of sequence motifs selected by Lembarg and Freeman (2007) characteristic for given rhomboid clades. The assignment to clades based only on presence of the characteristic motifs in proteins’ primary structure was, however, proven to be a good method of classification of rhomboid homologs since coincided well with phylogenetic analyses. The motif-based classification revealed that PARL-type proteases possess mostly 6 transmembrane domains and secretase B clade covers proteins with a divergent number of domains. The SB clade can however be further divided into 3 subclades (SB7, SB6 and SB3) containing proteins not only characterized by even numbers of transmembrane domains but primarily by the presence of a common arrangement of protein domains. SB7 consists of proteins containing only the rhomboid domain, while the SB6 and SB3 clades are constituted by proteins characterized by presence of UBA and zf-RanBP, respectively. The similar domain architecture within these 3 subclades suggests that proteins within these groups may share common functions.

The comparative genomic analysis also revealed that several inactive rhomboid-like proteins share high sequence identity with rhomboid considered as proteolytically active. These proteins formed common clusters with proteases belonging to different clades and contain, in their primary structure, motifs characteristic for given clades. Other inactive proteins clustered into two groups: Y-RL and Fx-RL. Proteins within these clusters share relatively high sequence identity, highly conserved primary structure motifs and domain architecture, which may indicate that they perform an important physiological function.

In most S2P proteases M50 domains are highly conserved, which is consistent with a previous study indicating the crucial role of some of these proteins in plant physiology and development. AraSP was proven to have a crucial role in plant development (Bötler et al. 2006) and AtEgy1 was shown to be involved in chloroplast biogenesis (Chen et al., 2005). The most conserved proteins contained, however, only a short C-terminal fragment of M50 domain and are probably proteolytically inactive. The importance and possible function of these proteins is hard to predict, since very few experimental data are available. The gene encoding AtEgy3 protein belonging to this group shows high coexpression with chaperonins and heat shock proteins involved, inter alia, in ER-associated protein degradation (Aoki et al. 2016).

Most sequences encoding aspartic intramembrane proteases were predicted to have nine transmembrane domains. We identified however several proteins which were predicted to have fewer transmembrane domains, but possess all features of active protease. Nevertheless, further studies are necessary to determine the actual number of the transmembrane domains as well as the proteolytic activity of these proteins. The most conserved group of proteins comprised homologs of AtSPP protein, which is consistent with previous data indicating that knockout mutation in the AtSPP gene is lethal (Hoshi et al. 2013). Also high evolutionary conservation of At SPPL1 homologs may indicate the key role of these proteins in plant growth and physiology. Surprisingly, the presenilin homologs seem to be the least conserved group among aspartic intramembrane proteins, but their function should not be underestimated, since they are found to participate in cytoskeletal related responses and chloroplast movement (Khandelwal et al. 2007).

## Founding

This work was supported by the National Science Center, Poland, based on decision number DEC-2014/15/B/NZ3/00412

## Literature

Aoki Y., Okamura Y., Tadak S., Kinoshita K., Obayashi T. (2016). ATTED-II in 2016: a plant coexpression database towards lineage-specific coexpression. Plant Cell Physiology 57, 1–9.

Bölter B., Nada A., Fulgosi H., Soll J. (2006). A chloroplastic inner envelope membrane protease is essential for plant development. FEBS Letters 580, 789–794.

Carpenter E.P., Beis K., Cameron A.D., Iwata S. (2008). Overcoming the challenges of membrane protein crystallography. Current Opinion in Structural Biology 18, 581–586.

Che P., Bussell J.D., Zhou W., Estavillo G.M., Pogson B.J., Smit S.M. (2010). Signaling from the Endoplasmic Reticulum Activates Brassinosteroid Signaling and Promotes Acclimation to Stress in Arabidopsis, Science Signaling 3, ra69.

Chen G., Bi Y.R., Li N. (2005). EGY1 encodes a membrane-associated and ATP-independent metalloprotease that is required for chloroplast development. Plant Journal 41, 364–375.

Chen G., Law K., Ho P., Zhang X., Li N. (2012). EGY2, a chloroplast membrane metalloprotease, plays a role in hypocotyl elongation in Arabidopsis, Molecular Biology Reports 39, 2147–2155.

Cowburn D. (1997). Peptide recognition by PTB and PDZ domains. Current Opinion in Structural Biology 7, 835–838.

Crooks G.E., Hon G., Chandonia J.M., Brenner S.E. (2004). WebLogo: A sequence logo generator. Genome Research 14, 1188–1190.

Dowse T.J., Soldati D., Soldati D. (2005). Rhomboid-like proteins in Apicomplexa: Phylogeny and nomenclature. Trends in Parasitology 21, 254–258.

Erez, E., Fass, D., Bibi E. (2009). How intramembrane proteases bury hydrolytic reactions in the membrane. Nature 459, 371–378. doi:10.1038/nature08146

Feng L., Yan H., Wu Z., Yan N., Wang Z., Jeffrey P.D., Shi Y. (2007). Structure of a Site-2 Protease Family Intramembrane Metalloprotease. Science 7, 1608–1612.

Feng L., Yan H., Wu Z., Yan N., Wang Z., Jeffrey P.D., Shi Y. (2007). Structure of a Site-2 Protease Family Intramembrane Metalloprotease. Science 318, 1608–1612.

Fluhrer, R. (2006). A γ-secretase-like intramembrane cleavage of TNF-αby the GxGD aspartyl protease SPPL2b, Nature Cell Biology 8, 894–896.

García-Lorenzo M., Sjödin A., Jansson S., Funk C. (2006). Protease gene families in Populus and Arabidopsis. BMC Plant Biology 6, 30 doi:10.1186/1471-2229-6-30.

Hoshi M., Ohki Y., Ito K., Tomita T., Iwatsubo T., Ishimaru, Y., et al. (2013). Experimental detection of proteolytic activity in a signal peptide peptidase of Arabidopsis thaliana. BMC Biochemistry 14, 16.

Kanaoka M.M., Urban S., Freeman M., Okada K. (2005). An Arabidopsis rhomboid homolog is an intramembrane protease in plants. FEBS Letters 579, 5723–5728.

Kanehisa M., Goto S. (2000). KEGG: kyoto encyclopedia of genes and genomes, Nucleic Acids Research 28, 27–30.

Khandelwal A., Chandu D., Roe C.M., Kopan R., Quatrano R.S. (2007) Moonlighting activity of presenilin in plants is independent of γ-secretase and evolutionarily conserved, Proceedings of the National Academy of Sciences 104, 13337–13342.

Knop M., Finger A., Braun T., Hellmuth K., Wolf D.H. (1996). Der1, a novel protein specifically required for endoplasmic reticulum degradation in yeast. The EMBO Journal 15, 753–763.

Lemberg, M. K., and Freeman, M. (2007). Functional and evolutionary implications of enhanced genomic analysis of rhomboid intramembrane proteases. Genome Res. 17, 1634–1646. doi.org/10.1101/gr.6425307

Lemberg, M. K., Menendez, J., Misik, A., Garcia, M., Koth, C. M., Freeman, M. (2005). Mechanism of intramembrane proteolysis investigated with purified rhomboid proteases. EMBO J. 24, 464–472.

Luo X., Hofmann K. (2001). The protease-associated domain: a homology domain associated with multiple classes of proteases. Trends in Biochemical Sciences 26, 147 – 148.

McQuibban G.A., Saurya S., Freeman M., Saurya S., Freeman M., Freeman M. (2003). Mitochondrial membrane remodelling regulated by a conserved rhomboid protease. Nature 423, 537–541.

Mueller T. D., Feigon J. (2002). Solution Structures of UBA Domains Reveal a Conserved Hydrophobic Surface for Protein–Protein Interactions. Journal of Molecular Biology 319, 1243–1255.

Ponting, C.P., Hutton M., Nyborg A., Baker M., Jansen K., Golde T.E. (2002). Identification of a novel family of presenilin homologues. Human Molecular Genetics 11, 1037–1044.

Ranganathan R., Ross E.M. (1997). PDZ domain proteins: Scaffolds for signaling complexes. Current Biology 7, R770–R773.

Rawlings, N.D., Barrett, A.J., Finn, R.D. (2016). Twenty years of the MEROPS database of proteolytic enzymes, their substrates and inhibitors. Nucleic Acids Res 44, 343–350.

Rawson R.B., Zelenski N.G., Nijhawan D., Ye J., Sakai J., Hasan M.T., et al. (1997). Complementation Cloning of S2P, a Gene Encoding a Putative Metalloprotease Required for Intramembrane Cleavage of SREBPs. Mol. Cell 1, 47–57.

Schuhmann H., Huesgen P.F., Adamska I. (2012). The family of Deg/HtrA proteases in plants. BMC Plant Biology 12, 52.

Schwacke R., Schneider A., Van Der Graaff E., Fischer K., Catoni E., Desimone M., Frommer W.B., Flügge U.I., Kunze R. (2003). ARAMEMNON, a Novel Database for Arabidopsis Integral Membrane Proteins. Plant Physiology 131, 16–26.

Simossis, V.A., Heringa, J. (2005). PRALINE: a multiple sequence alignment toolbox that integrates homology-extended and secondary structure information. Nucleic Acids Research 33, W289–W294. http://doi.org/10.1093/nar/gki390]

Steggerda S.M., Paschal B.M. (2002). Regulation of nuclear import and export by the GTPase Ran. International Review of Cytology 217, 41–91.

Tanya Z., Berardini, L.R., Li D., Mezheritsky Y., Muller R., Strait E., Huala E. (2015). The Arabidopsis Information Resource: Making and mining the “gold standard” annotated reference plant genome. Genesis, doi: 10.1002/dvg.22877.

Tomita T., Watabiki T., Takikawa R., Morohashi Y., Takasugi N., Kopan R., de Strooper B., Iwatsubo, T. (2001). The first proline of PALP motif at the C terminus of presenilins is obligatory for stabilization, complex formation and γ-secretase activities of presenilins. Journal of Biological Chemistry 276, 33273–33281.

Tripathi L.P., Sowdhamini R. (2006). Cross genome comparisons of serine proteases in Arabidopsis and rice. BMC Genomics 7, 200.

Wang J., Beher, D., Nyborg A. C., Shearman M. S., Golde T.E., Goate, A. (2006). C-terminal PAL motif of presenilin and presenilin homologues required for normal active site conformation. Journal of Neurochemistry 96, 218–227. doi:10.1111/j.1471-4159.2005.03548.

Yaseen N.R., Blobel G. (1999). Two distinct classes of Ran-binding sites on the nucleoporin Nup-358. Proceedings of The National Academy of Science U.S.A. 96, 5516–5521.

Zybailov B., Rutschow H., Friso G., Rudella A., Emanuelsson O., Sun Q., van Wijk K.J. (2008). Sorting signals, N-terminal modifications and abundance of the chloroplast proteome, PLoS One 23, e1994.

